# Assessing extracellular vesicle proteins as predictive biomarkers for developing type 1 diabetes

**DOI:** 10.64898/2026.02.06.703600

**Authors:** Panshak P. Dakup, Lisa M. Bramer, Athena Schepmoes, Ivo Diaz Ludovico, Javier Flores, Raghavendra G. Mirmira, Bobbie-Jo M. Webb-Robertson, Thomas O. Metz, Emily K. Sims, Ernesto S. Nakayasu

## Abstract

Plasma extracellular vesicles (EVs) are considered excellent sources for biomarker discovery since they carry signatures of their cellular origin and disease processes. In this paper, we evaluate the potential of plasma EV proteomics analysis for identifying predictive biomarkers of developing type 1 diabetes (T1D), which results from autoimmune destruction of insulin-producing β cells in the islet. We used strong anion exchange beads (Mag-Net) to capture plasma EVs from 19 donors with islet autoimmunity (diagnosed by circulating autoantibodies against islet proteins – AAB+) vs. 17 control individuals and analyzed their protein cargo by mass spectrometry. The analysis identified and quantified 5,480 proteins, a 3.2-fold increase in proteome coverage compared to our previous T1D biomarker proteomics study that used whole plasma depleted of the 14 most abundant proteins. The Mag-Net approach also detected 1,306 out of the 1,717 proteins (76%) that we previously verified as EV proteins. Statistical tests revealed 448 proteins to be differentially abundant in AAB+ vs control volunteers, including 69 previously verified EV proteins. A functional-enrichment analysis resulted in overrepresentation of 25 pathways among the differentially abundant proteins, including pathways related to autoimmune response and lipid metabolism. The capacity of this data to predict AAB+ was tested with a machine learning analysis using a random forest model, resulting in a receiver operating characteristic-area under the curve of 0.81. Overall, our study indicates that plasma EV proteomics analysis can be an exciting approach for studying biomarkers for developing T1D.

**Significance of the study:** Type 1 diabetes (T1D) is a disease characterized by the body’s inability to produce insulin and consequently, to control blood glucose levels. Despite the initial trigger being unclear, the disease development process involves an autoimmune response to the islets of Langerhans, resulting in the death of insulin-producing β cells. There is no cure for the disease, and treatment relies on exogenous administration of insulin. Therefore, preventive therapies that block the autoimmune process are attractive for treating T1D. In fact, anti-CD3 antibody (Teplizumab) delays the onset of T1D by 2 years by targeting T cells. Predictive biomarkers for developing T1D are needed to aid the development and implementation of new therapies and to identify the initial trigger and mechanisms of the islet autoimmune process. In this paper, we assess the potential of plasma extracellular vesicle (EV) proteomics analysis for identifying predictive biomarkers of T1D. Our results show excellent potential of the approach, opening opportunities to perform broader studies to identify biomarkers for developing T1D.

## Main Text

Cells secrete lipid-bilayer delimited extracellular vesicles (EVs) carrying a variety of biomolecules, including proteins, lipids, metabolites and nucleic acids [1]. EVs play import roles in inter-cellular communication and are classified mainly as exosomes and ectosomes, based on their biogenesis [1]. Exosomal biogenesis involves invagination of the membranes from organelles named multivesicular body, resulting in 30-100 nm diameter vesicles. These EVs are released into the extracellular environment when the multivesicular body membrane fuses with the plasma membrane. Some classical exosome markers include the tetraspanins CD9, CD63 and CD81 [1]. In an alternative biogenesis mechanism, EVs are budded directly from plasma membranes, releasing ectosomes. There are a variety of ectosome subtypes, but perhaps the best known are the microvesicles, which are 100-1000 nm in diameter and carry markers such as CD40 [1]. EVs represent signatures of their cell of origin and associated disease processes, making them excellent sources for biomarker development [2-6]. However, biofluids that contain EVs, such as blood plasma, are highly complex and have proteins and particles with similar physicochemical properties, a challenge for obtaining preparations with purity and quantity required for molecular analyses [7, 8].

Considering this challenge, we previously conducted a meta-analysis of public proteomics data to establish a verified protein composition within EVs [9], allowing us to then focus experimental efforts less on the purity of the EV preparation [10] and more on sensitive quantification in small sample sizes [11]. Traditional EV purification methods often require 1-2 mL of plasma for proteomics analysis, which can be prohibitive for large clinical studies in which multiple analyses are performed within the available samples. To address this challenge, we exploited the principle that smaller diameter size-exclusion chromatography columns require less material and performed purifications on a column with 5 mm inner diameter, reducing the sample requirement to only 50 μL of plasma. This approach resulted in the identification of 838 proteins [11]. More recently, Wu et al. developed a fully automated method for enriching plasma EVs based on strong-anion exchange magnetic beads (Mag-Net), leading to the identification and quantification of over 4000 proteins [12]. Likewise, this method requires only 50 μL of plasma.

Our biomarker discovery studies have focused on the autoimmune disease type 1 diabetes (T1D). T1D results from the autoimmune destruction of pancreatic β cells, leading to the body’s inability to control blood glucose levels [13]. This process is preceded by immune cell infiltration in the islets of Langerhans, which can be diagnosed by appearance of circulating autoantibodies against islet proteins (seroconversion) [13]. Once β-cell mass is lost, there are no effective therapies to regenerate these cells. Therefore, many efforts have concentrated on developing therapies that can prevent β-cell loss, including FDA-approved anti-CD3 monoclonal antibodies that can delay the disease onset by 2 years [14, 15]. In this context, biomarkers that precisely predict disease development are needed to guide accurate implementation of preventive therapies. We previously conducted a proteomics study of The Environmental Determinants of Diabetes in the Young (TEDDY) consortium and identified peptide panels that predict 6 months prior to seroconversion if a child will develop T1D (receiver operating characteristic area under the curve - ROC-AUC=0.92) or only islet autoimmunity (ROC-AUC=0.87) by the age of 6 years [16]. One of these biomarkers, platelet basic protein (PPBP/CXCL7), is indeed an EV protein that regulates pancreatic β-cell apoptosis and macrophage activity [9, 16]. PD-L1 is another biomarker candidate, as its abundance was elevated in plasma EVs from islet autoantibody-positive individuals [17]. Moreover, proteomics analysis of EV-enriched plasma from non-obese diabetic mouse models identified proteins related to β-cell function and immune regulation, indicating that EVs may carry signatures of disease pathogenesis [18]. These studies support the potential of EVs as a source for identifying novel biomarkers for developing T1D.

In addition to the potential for EVs as a source for identifying tissue-specific protein biomarkers, they also represent an excellent opportunity to improve plasma proteome coverage during mass spectrometry (MS)-based proteomics analysis. MS-based analysis of total plasma proteins leads to the detection of only approximately 600 to 1000 proteins due to bias towards highly abundant plasma proteins [12]. Depletion of the top ∼15 most abundant plasma proteins improves the proteome coverage to 1,500-2,000 proteins [16, 19]. As an alternative strategy, analyzing plasma EVs can further improve coverage of the proteome to >4,000 proteins [12]. Here, we assess the potential of plasma EV for identifying biomarkers for developing T1D. We obtained plasma from autoantibody positive (AAB+) and control (autoantibody negative) volunteers and submitted these to Mag-Net EV enrichment and preparation for proteomics analysis (**Fig. 1 and Tab. 1**). The resulting peptides were analyzed using data-independent acquisition analysis on an Astral mass spectrometer, and a total of 5480 proteins were identified and quantified (**Tab. S1**). The analysis identified 1306 (76%) out of the 1717 proteins that were verified as being from EVs in our previous proteomics meta-analysis (**Fig. 2A**) [9]. Moreover, this analysis increased by 3.2-fold the coverage of the plasma proteome compared to our previous T1D biomarker study [16], including the detection of 1,391 (81%) of the 1,720 found in that study (**Fig. 2B**). Overall, these results showed that the Mag-Net approach can deepen the plasma proteome coverage and detect a large portion of the verified EV proteins identified in our previous meta-analysis of public proteomics data.

**Figure 1:**
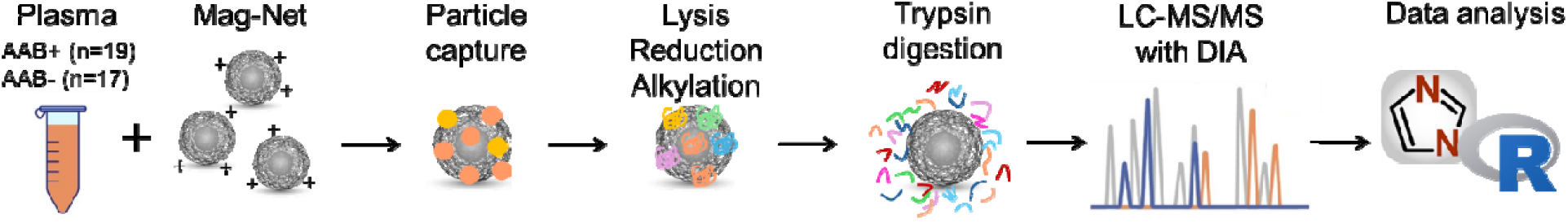
Workflow of EV enrichment using Mag-Net beads

**Figure 2:**
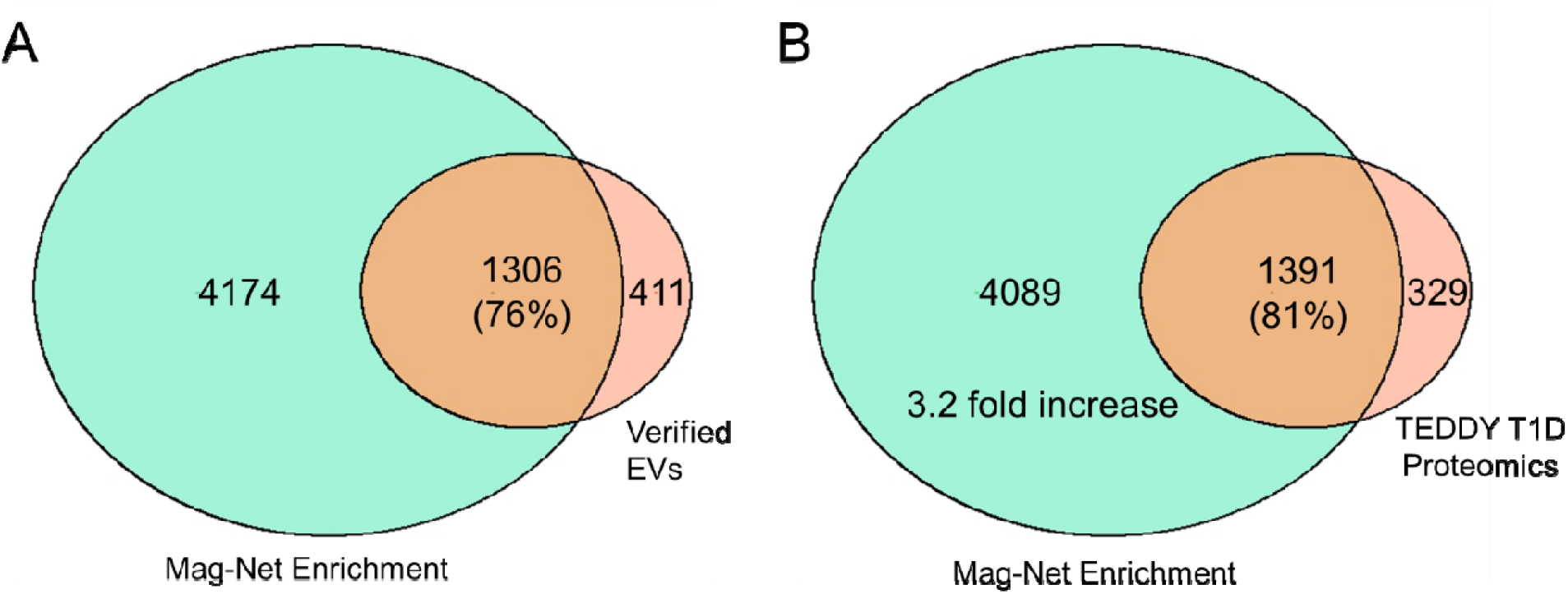
Proteomic coverage achieved with the Mag-Net Enrichment Approach. Venn diagrams show A) the number/percentage of proteins identified by the Mag-Net enrichment compared to verified EV proteins identified in the meta-analysis by Vallejo et al [9]. B) the increase in depth of plasma proteome coverage with Mag-Net enrichment compared to immunodepletion and fractionation-based proteomics in the TEDDY study [16].

We next aimed to identify proteins that were differentially abundant between AAB+ and control groups. During quality control analysis of the data, one sample in the control group was identified as a potential outlier based on the robust Mahalanobis distance-based algorithm, which was further confirmed to be a statistical outlier via principal component analysis (PCA) and thus removed. This resulted in a final comparison of 19 samples from AAB+ donors vs. 17 samples from control individuals (**Tab. 1**). A statistical analysis comprised of ANOVA and G-test identified 448 proteins to be differentially abundant in AAB+ vs control volunteers (p≤0.05 for either test), with 100 and 348 proteins higher and lower in AAB+, respectively (**Fig. 3A**). Furthermore, 69 significant proteins were verified as EV proteins in our previous meta-analysis, including 32 and 37 proteins found to be higher and lower in AAB+, respectively [9]. Among the most regulated proteins, YPEL5, NXN and MMP3 were downregulated, and RARRES1 and CBLN2 were upregulated (**Fig. 3A**). Among the previously verified EV proteins, PEPD and APOBEC3C were among the most downregulated proteins, whereas CD14 and HBD were among the most upregulated proteins (**Fig. 3A**). A functional-enrichment analysis of the differentially abundant proteins showed enrichment in 25 pathways by Fisher’s exact test with a p≤0.05 cutoff (**Fig. 3B**). This includes pathways involved in autoimmune diseases (“Neutrophil extracellular trap formation” and “Systemic lupus erythematosus”) and lipid metabolism (“Alcoholism”, “Fatty acid metabolism”, “Glycerophospholipid metabolism” and “Fatty acid biosynthesis”) (**Fig. 3B**). Out of the 25 pathways, 11 (44%) were previously shown to be also enriched among the verified EV proteins (**Fig. 3B**) [9]. Together these results identified biomarker candidates of islet autoimmunity coming from a variety of pathways, with a substantial portion of these proteins previously verified to be EV proteins.

**Figure 3:**
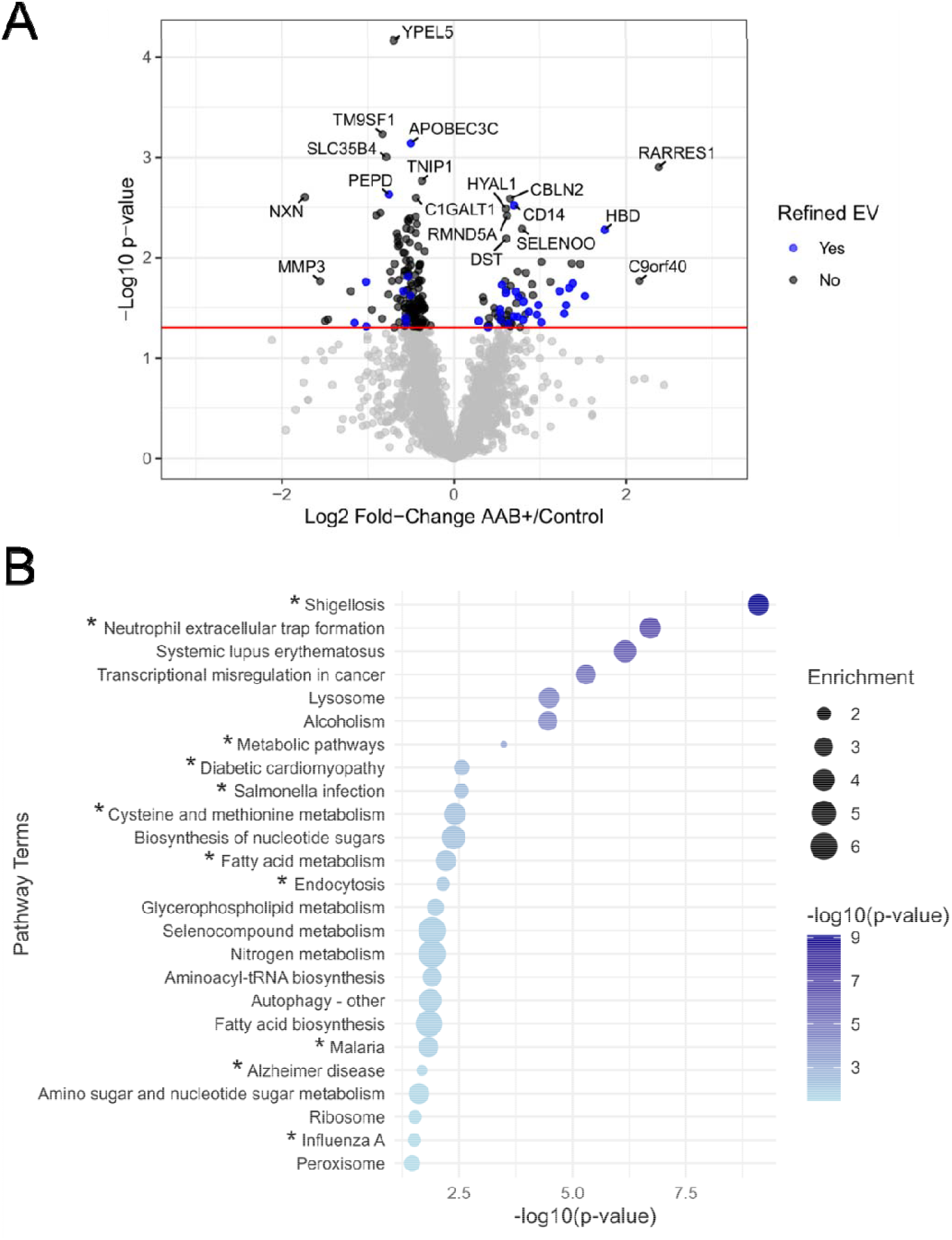
Differential analysis of identified proteins. A) Volcano plot showing ANOVA statistical results, where the red horizonal line corresponds to a p-value of 0.05. Proteins previously verified as EVs by Vallejo et al [9] are shown in blue. B) Bubble plot summary of the significant pathways identified by DAVID functional-enrichment analysis. The y-axis represents the pathway terms while the x-axis displays the Fisher’s exact test p-values (-log10(p-value)). Bubble size corresponds to the fold enrichment and color gradient reflects significance of each pathway. Asterisks (*) indicate pathways that were also enriched in verified EV proteins.

To further assess the data, we performed a machine learning analysis to identify panels of proteins that predict autoantibody positivity. We used a random forest model to fit the data and constructed a receiver operating characteristic (ROC) curve, which showed an area under the curve (AUC) of 0.81 (**Fig. 4**). This model was based on 421 proteins, including the 63 proteins that were previously verified as EV proteins in our proteomics meta-analysis (**Tab. S1**). These results demonstrate the potential of plasma EV proteomics analysis for identifying biomarkers for developing T1D.

**Figure 4.**
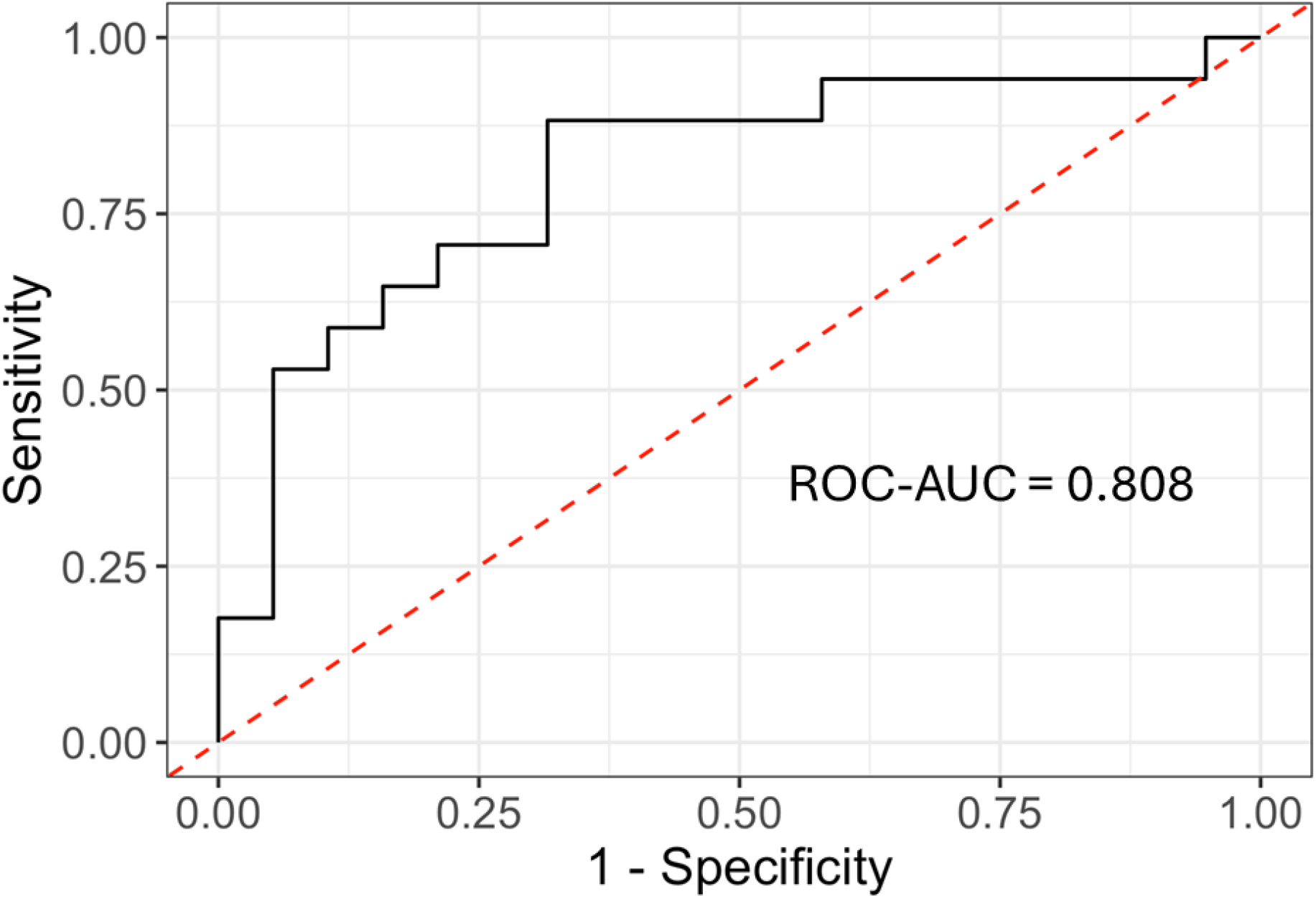
Prediction of islet autoimmunity by multivariate panel machine learning analysis. The graph shows a receiver operating characteristic (ROC) curve of predicting autoantibody positivity (AAB+, islet autoimmunity) vs. control.

Overall, here we showed that the analysis of EVs can be an excellent alternative approach to develop biomarkers for T1D, compared with conventional proteomics analysis of whole or immunodepleted plasma. The application of the Mag-Net approach to enrich for EVs increased the coverage of the plasma proteomics coverage by 3.2-fold compared to our previous analysis [16]. A recent independent study compared multiple methods for plasma proteomics analysis, indicating additional promising alternative approaches, such as SEER Proteograph XT and PreOmics ENRICHplus, although these do not enrich EVs [20]. Therefore, they might not bring the added benefit of information on the tissue of origin. Our analysis also led to the identification of potential biomarker candidates, which include proteins from a variety of functions. These pathways include those related to autoimmune response, such as neutrophil extracellular trap formation. Neutrophil extracellular traps have been detected in plasma from individuals recently diagnosed with T1D using immunoassays [21]. Several pathways related to lipid metabolism were also found to be regulated in AAB+ individuals in our analysis. In fact, the plasma lipid composition is altered during T1D development [22, 23]. We have found that free fatty acids are increased in plasma in AAB+ individuals and that such free fatty acids can enhance the killing of β cells [24]. Therefore, in addition to biomarkers, the molecular signatures found in biomarker development analysis can also provide mechanistic insights into disease development. Lastly, the machine learning analysis identified a multivariate panel that can predict autoantibody positivity with an ROC AUC of 0.81, further supporting the potential of targeting EVs for T1D biomarker development.

### Limitations of the study

This is an initial feasibility study to assess the potential of targeting EVs for biomarkers of developing T1D. Biomarker development requires multiple phases: discovery, verification (sometimes referred as validation) and validation (also referred to as clinical and analytical validation) [25]. Therefore, the biomarkers identified in this paper should be considered only candidates and will need additional rounds of evaluation before they can be employed in the clinic.

## EXPERIMENTAL PROCEDURES

### Blood plasma samples

Plasma samples were obtained from the Indiana University (IU) Center for Diabetes and Metabolic Diseases biobank. The protocol was approved by the IU Institutional Review Board, and informed consent following the Declaration of Helsinki guidance was obtained. Samples of autoantibody positive individuals were matched to controls based on age, sex and BMI (**Table 1**), and shipped deidentified for mass spectrometry analysis. Samples were randomized by the team statistician and laboratory staff that prepared and analyzed the samples had no access to the metadata information.

**Table 1.** Characteristics of the donors.

|  | AAB+ | Control | p-value |
| --- | --- | --- | --- |
| # Donors | 19 | 17 |  |
| Age | 14.2 ± 9.7 | 16.3 ± 12.0 | 0.57 |
| Sex | M: 15 (79%)<br>F: 4 (21%) | M: 14 (79%)<br>F: 3 (21%) | 1 |
| BMI | 19.7 ± 5.8 | 19.9 ± 4.2 | 0.66 |

### Mag-Net EV enrichment

Plasma EVs were enriched using a magnetic affinity–based protocol adapted from Wu et al. [12], employing size- and charge-based capture on strong anion exchange beads with automated handling on a Thermo Scientific KingFisher Flex. MagReSyn strong anion exchange beads (12.5 µL per sample) were equilibrated twice with 200 µL wash buffer (50 mM BisTrispropane, pH 6.3, 150 mM NaCl). Plasma (50 µL) was mixed 1:1 with binding buffer (100 mM BisTrispropane, pH 6.3, 150 mM NaCl) supplemented with EDTA-free Pierce Protease Inhibitor Cocktail. The diluted plasma was added to the equilibrated SAX beads and incubated for 30 min with mixing. Beads were then washed three times with wash buffer for 5 min each. EV enriched particles bound to the beads were solubilized and reduced in 100 µL lysis/reduction buffer (50 mM Tris-HCl, pH 8.5, 1% sodium dodecyl sulfate, 10 mM Tris-(2-carboxyethyl)phosphine), followed by alkylation with 15 mM iodoacetamide for 30 min in the dark at room temperature. Proteins were precipitated by addition of acetonitrile to a final concentration of 70% (v/v), and beads were washed three times with 95% acetonitrile followed by three washes with 70% ethanol. On-bead digestion was performed in 25 mM ammonium bicarbonate with sequencing grade trypsin at an enzyme-to-protein ratio of 1:20 (w/w) at 47°C for 2 h with mixing. Digestion was quenched by adding trifluoroacetic acid to a final concentration of 0.5% (v/v). Peptides were desalted using a Waters Oasis HLB µElution plate and eluted with 70:30 acetonitrile:water (v/v). Peptide concentrations were determined by BCA assay, and samples were stored at −80°C until LCMS/MS analysis.

### Mass spectrometry analysis

For each analysis, 500 ng of peptides were injected into a Thermo Scientific Orbitrap Astral mass spectrometer coupled to a Thermo Scientific Vanquish Neo UHPLC system. Chromatographic separation was performed on an ES 906 PepMap C18 column (150 mm × 150 µm i.d., 2 µm, 100 Å) at a flow rate of 3.5 µL/min. Mobile phase A was 0.1% formic acid in water, and mobile phase B was 0.1% formic acid in acetonitrile. Peptides were separated using a 22-min gradient, followed by an 8 min high organic wash and column re-equilibration at initial conditions. Data were acquired in data-independent acquisition (DIA) mode. For each injection, MS1 spectra were acquired over 400–1000 m/z using fixed 2 m/z DIA isolation windows. MS2 spectra were acquired over 150–2000 m/z. The Orbitrap resolution for MS1 was set to 240,000 with a normalized AGC target of 500%. RAW files were processed in DIA-NN 2.2.0 in library-free mode with *in-silico* library generation [26]. Spectra were searched against tryptic peptides (up to two missed cleavages, peptide length 7–30 amino acids). Carbamidomethylation of cysteine was set as a fixed modification, and variable modifications included methionine oxidation and protein N-terminal acetylation (up to three variable modifications per peptide). Peptide and protein level false-discovery rates were controlled at 1% (q < 0.01).

### Statistical analysis

All statistical analyses were conducted in R version 4.4.3 using the *pmartR* package [27, 28]. Data was transformed into log2 scale and missing observations were denoted with NA values. Proteins without sufficient data to conduct quantitative or qualitative statistics were filtered from the data [29]. The data was evaluated for outlier samples using a robust Mahalanobis distance-based algorithm using the recommended p-value threshold of 0.0001 [30]. Data was normalized using median centering. For each protein, an analysis of variance (ANOVA) model was fit to the data and tested for a difference in mean relative abundances for the AAB+ and control groups.

Additionally, a G-test was performed on each protein to test for a difference in the mean probability of a protein being detected in the AAB+ and control groups [30]. Functional-enrichment analysis was performed with DAVID, using the KEGG annotation [31]. Machine learning was conducted to evaluate the ability to differentiate AAB+ and control samples in a predictive manner. Feature screening was first conducted, retaining proteins with an ANOVA or G-test p-value of 0.1 or less as potential predictors. Potential predictors without sufficient data to conduct an ANOVA were transformed to binary values corresponding to the protein being detected or not in a sample. All other potential predictors were used quantitatively, and missing values were imputed using expectation maximization (EM) imputation [32]. A random forest model [33] was fit to the data, using out-of-bag samples for validation, and the area under the curve (AUC) was computed.

## ACKNOWLEDGMENTS

Part of the work was performed in the Environmental Molecular Sciences Laboratory, a U.S. Department of Energy (DOE) national scientific user facility at Pacific Northwest National Laboratory (PNNL) in Richland, WA. Battelle operates PNNL for the DOE under contract DE-AC05-76RLO01830. This work was supported by the National Institute of Diabetes and Digestive and Kidney Diseases grants R01 DK138335 (to E.S.N., B-J.M.W-R., and T.O.M.), R01 DK133881 (to E.K.S. and R.G.M.), U01 DK127786 (to R.G.M. and E.S.N.), and R01 DK060581 (to R.G.M.). E.S.N. was also supported by the Human Islet Research Network Catalyst Award (via U24 DK104162).

## SUPPORTING INFORMATION

Supplemental table S1 is available online.

